# *De novo* design and structure of a peptide-centric TCR mimic binding module

**DOI:** 10.1101/2024.12.16.628822

**Authors:** Karsten D. Householder, Xinyu Xiang, Kevin M. Jude, Arthur Deng, Matthias Obenaus, Steven C. Wilson, Xiaojing Chen, Nan Wang, K. Christopher Garcia

## Abstract

T cell receptor (TCR) mimics offer a promising platform for tumor-specific targeting of peptide-MHC in cancer immunotherapy. Here, we designed a *de novo* α-helical TCR mimic (TCRm) specific for the NY-ESO-1 peptide presented by HLA-A*02, achieving high on-target specificity with nanomolar affinity (K_d_ = 9.5 nM). The structure of the TCRm/pMHC complex at 2.05 Å resolution revealed a rigid TCR-like docking mode with an unusual degree of focus on the up-facing NY-ESO-1 side chains, suggesting the potential for reduced off-target reactivity. Indeed, a structure-informed *in silico* screen of 14,363 HLA-A*02 peptides correctly predicted two off-target peptides, yet our TCRm maintained a wide therapeutic window as a T cell engager. These results represent a path for precision targeting of tumor antigens with peptide-focused α-helical TCR mimics.

## Main Text

Targeting peptide-major histocompatibility complex (pMHC) molecules on tumor cells with high specificity is a long-standing goal in cancer immunotherapy (*1, 2*). Class I MHC molecules present short peptide fragments (8-13 amino acids) from the intracellular proteome on the cell surface, providing a snapshot of the internal state of the cell (ie – healthy, cancerous, infected, etc.) (*3–5*). Cancer cells present mutant or self-peptides that are recognized by T cell receptors (TCRs) through a mechanism known as ‘MHC restriction’, which interacts with both the MHC and the peptide composite surface (*6*). This recognition enables T cells to selectively detect and eliminate cancerous cells.

Therapeutic application of natural TCRs is hindered by their low affinity (*7–9*). To overcome these limitations, many groups have engineered TCR mimic (TCRm) antibodies – high affinity molecules that replicate the MHC-restricted peptide recognition of TCRs (*10–17*). These mimics are particularly valuable as cancer-specific probes and therapeutics. A recent example is tebentafusp-tebn, a bispecific T cell engager (BiTE) approved in 2022 for uveal melanoma (*18– 19*). Despite these advantages, the discovery of TCR mimics remains a significant technical challenge. Antibodies have not been naturally selected for pMHC restriction, like TCRs, and generally exhibit interactions with the MHC helices that result in substantial potential for recognizing peptide-MHC from normal tissue, leading to clinical toxicities (*20–24*). Although strategies such as “repurposing” TCRm antibodies have been described (*10, 25*), fine-tuning these antibodies to exhibit acceptably “clean” peptide specificity represents a major obstacle for the field (*20, 26–28*).

To address these limitations, we sought to develop α-helical TCR mimics that could be rapidly programmed into cancer-specific T cell engagers. α-helical bundles offer several advantages over antibodies: they are compact, modular, and present amino acid side chains on a relatively rigid platform for interactions with ligands (*29*), reducing the design complexity observed with flexible complementarity-determining region (CDR) loops on antibodies. Furthermore, CDR loop flexibility is a mechanism of cross-reactivity by TCRs (*30–32*), so more rigid scaffolds could minimize structural adaptation to alternative peptides. Additionally, state-of-the-art protein design tools like RFdiffusion and ProteinMPNN (*33, 34*), which have demonstrated increased success rates for *de novo* protein design, exhibit a natural bias towards α-helical folds.

We show that an integrated pipeline of computational design, high-throughput screening, and structural validation, can streamline the discovery of highly peptide-specific TCR mimics essentially “out of the box.” Our approach leverages RFdiffusion to generate α-helical scaffolds, ProteinMPNN to design sequences, yeast display for on-target selections, and X-ray crystallography for structural validation. The resulting high-resolution structures confirm on-target binding but also inform off-target specificity and optimization strategies.

### Design of α-helical “mini”-TCR mimics with RFdiffusion and ProteinMPNN

We targeted the NY-ESO-1_157-165 (C9V)_ peptide presented by HLA-A*02, a well-characterized tumor antigen (*2, 26, 35*). We sought to generate four-helix α-helical bundles, which are thermostable and found across the natural proteome to mediate diverse protein-protein interactions. Helical bundles, such as cytokines, are also ideal engineering scaffolds with proven robustness to amino acid substitutions on the helical faces (*36*). The protein design tools RFdiffusion and ProteinMPNN have also been found to bias towards α-helical folds, so we reasoned we might achieve higher experimental success rates by restricting to this fold space. First, we generated 50 monomer folds with RFdiffusion ranging between 80-100 amino acids and visually inspected the results for symmetrical and tightly packed bundles with a high likelihood of expression as recombinant proteins. We selected the best such fold (99 amino acids) to be used as a template scaffold for RFdiffusion fold-conditioning over the peptide-MHC target. We selected the NY-ESO-1_157-165 (C9V)_ peptide presented on the HLA-A*02 allele as our model antigen due to existing structures in the Protein Data Bank (PDB) and robust experimental systems for downstream validation. AlphaFold2 was then used to predict the structure of the NY-ESO-1 HLA-A*02 β2M input (*37*). We next specified the up-facing Met4, Trp5, Thr7, and Gln8 peptide residues as hotspots for design and masked the loops of our input scaffold to generate additional structural diversity at inference time. We produced 100 unique four-helix bundles docked over NY-ESO-1 HLA-A*02, and for each one, sampled 6 sequences with ProteinMPNN for 600 designs total. Each design complexed with our target was then predicted by AlphaFold2 and sorted by the interface predicted aligned error (iPAE) metric, as previously described (*38*). 4 out of 100 scaffolds yielded at least one design under an iPAE threshold of 10.0 and therefore, was considered a “hit” scaffold (Fig. 1A).

**Fig. 1.**
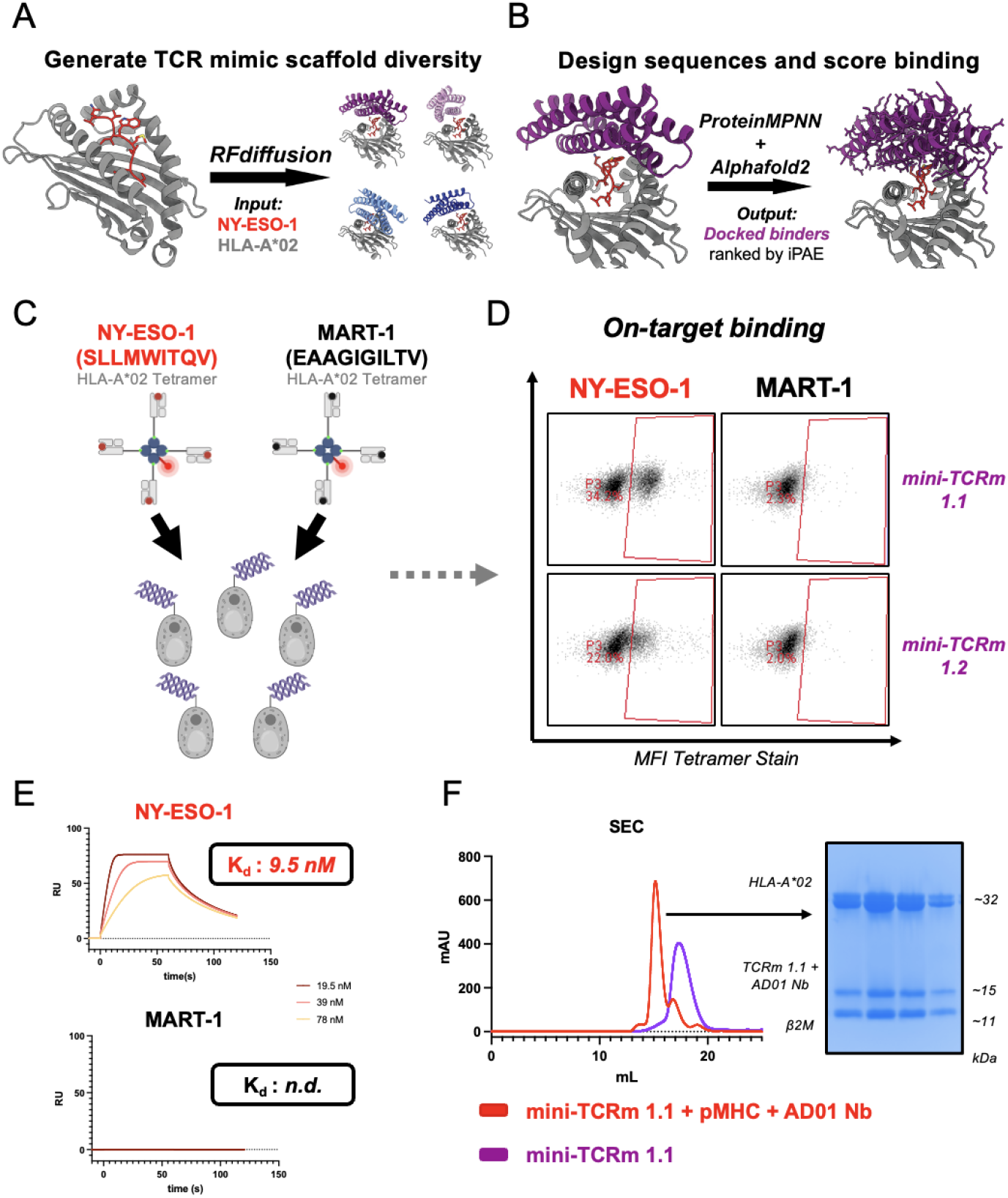
Design and experimental validation of *de novo* mini-TCR mimics. **(A)** Schematic of RFdiffusion fold-conditioning pipeline to generate four scaffolds over the NY-ESO-1 HLA-A*02 input (red and gray). **(B)** Schematic of pipeline using ProteinMPNN to design sequences for Scaffold #1 (purple) and predict binders with AlphaFold2. **(C)** Yeast display screening of top *de novo* designs with NY-ESO-1 (red) vs. MART-1 HLA-A*02 (black) tetramers. **(D)** Binders staining NY-ESO-1 but not MART-1 HLA-A*02 tetramer by flow cytometry. **(E)** SPR analysis of mini-TCRm 1.1 analyte with immobilized NY-ESO-1 HLA-A*02 (red) vs. MART-1 HLA-A*02 (black). Dissociation constants indicated on sensograms (K_d_); n.d., not determined; RU, response unit. **(F)** SEC profiles of mini-TCRm 1.1 (purple) vs. co-eluting complex (red) of mini-TCRm 1.1, NY-ESO-1 HLA-A*02, and AD01 nanobody. SDS-Page gel showing fractions of co-eluting complex pooled for crystal screens. mAU, milli-Absorbance Units; kDa, kilodaltons.

Our top scaffolds varied in helical bundle arrangement and size (99-129 amino acids). The docking orientation over the peptide-MHC complex also differed for each predicted binder, with some anticipated to make more MHC versus peptide contacts than others. To explore the sequence space of each hit more deeply, we generated 500 sequences per hit scaffold with ProteinMPNN, for 2000 designs total, and again predicted and scored each complex with AlphaFold2. Scaffold #1 yielded the highest computational success rates, with 129/500 designs passing the iPAE threshold (Fig. 1B). All other scaffolds yielded hits, but at lower frequencies than Scaffold #1. Furthermore, upon visual inspection, we noticed several desirable properties of Scaffold #1 versus the other scaffolds. First, it exhibited a centered, TCR-like docking mode over the peptide-MHC target, making it more likely to produce peptide-specific contacts. Second, all four specified hotspot residues were predicted contacts in several of the top designs. Taken together, Scaffold #1 appeared to best meet our design criteria, so we proceeded with the lowest iPAE-scoring binders for screening.

### Experimental validation of *de novo* TCR mimics specific to NY-ESO-1 HLA-A*02

The top five designs were displayed as single clones on yeast cells, and then stained with 100 nanomolar NY-ESO-1 or MART-1 HLA-A*02 tetramer (Fig. 1C). The MART-1 tetramer served as a negative control to ensure binders were NY-ESO-1-specific. Indeed, we found 2 out of 5 designs were FACS stained by the NY-ESO-1 tetramer but not MART-1 (Fig. 1D). We selected the strongest NY-ESO-1 binder (mini-TCRm 1.1) for recombinant expression in BL21 *E. Coli* and purified by Nickel-NTA and size-exclusion chromatography (SEC). The recombinant protein was soluble with high yields and eluted as a single peak by SEC (see mini-TCRm 1.1 peak in Fig. 1F). We then performed surface plasmon resonance (SPR) with this binder to interrogate its binding kinetics and affinity to NY-ESO-1 versus MART-1 HLA-A*02. SPR analysis revealed a dissociation constant (K_d_) of 9.5 nM for NY-ESO-1 HLA-A*02, but no detectable affinity for MART-1 HLA-A*02 (Fig. 1E). This indicated that mini-TCRm 1.1 is peptide-specific, with undetectable trace affinity for the same MHC presenting a different peptide. Finally, we complexed the binder with refolded NY-ESO-1 HLA-A*02 and the AD01 anti-beta-2-microglobulin (β2M) nanobody, which served as a crystallization chaperone. This complex co-eluted by SEC (Fig. 1F) and readily crystallized during screening.

### High-resolution crystal structure reveals peptide-specific interactions

We determined the crystal structure of the complex at 2.05 Å resolution (Fig. 2, A and B; table S1). The binder adopts a diagonal TCR-like docking footprint at a 40-degree angle relative to the peptide-MHC groove (Fig. 2C). The total buried surface area at the mini-TCRm/pMHC interface is 1216.7 Å^2^, with approximately 31% contribution from peptide-specific contacts. The mini-TCR mimic uses 14 residues within the groove of the A2 and A3 helices to form a concentrated network of hydrophobic and polar peptide contacts. Strikingly, the peptide-contacting side chains appear to bend inward creating a shape-complementary shell around the peptide side chains. In particular, the Met4-Trp5 motif of NY-ESO-1 bulges prominently above the MHC groove and inserts into two hydrophobic pockets between the A2 and A3 helices (Fig. 2, B and D), stabilized by three hydrogen bonds (Fig. 2E). The binder also evenly disperses hydrophobic contacts across both helices of HLA-A*02. These are supplemented by five hydrogen bonds and one salt bridge to the MHC, helping orient the binder precisely over the peptide (Fig. 2F). Notably, three of these hydrogen bonds (formed by Asn54, Asn80, and Arg57 of the binder) interact with residues inside the HLA-A*\**02 groove around Trp5.

**Fig. 2.**
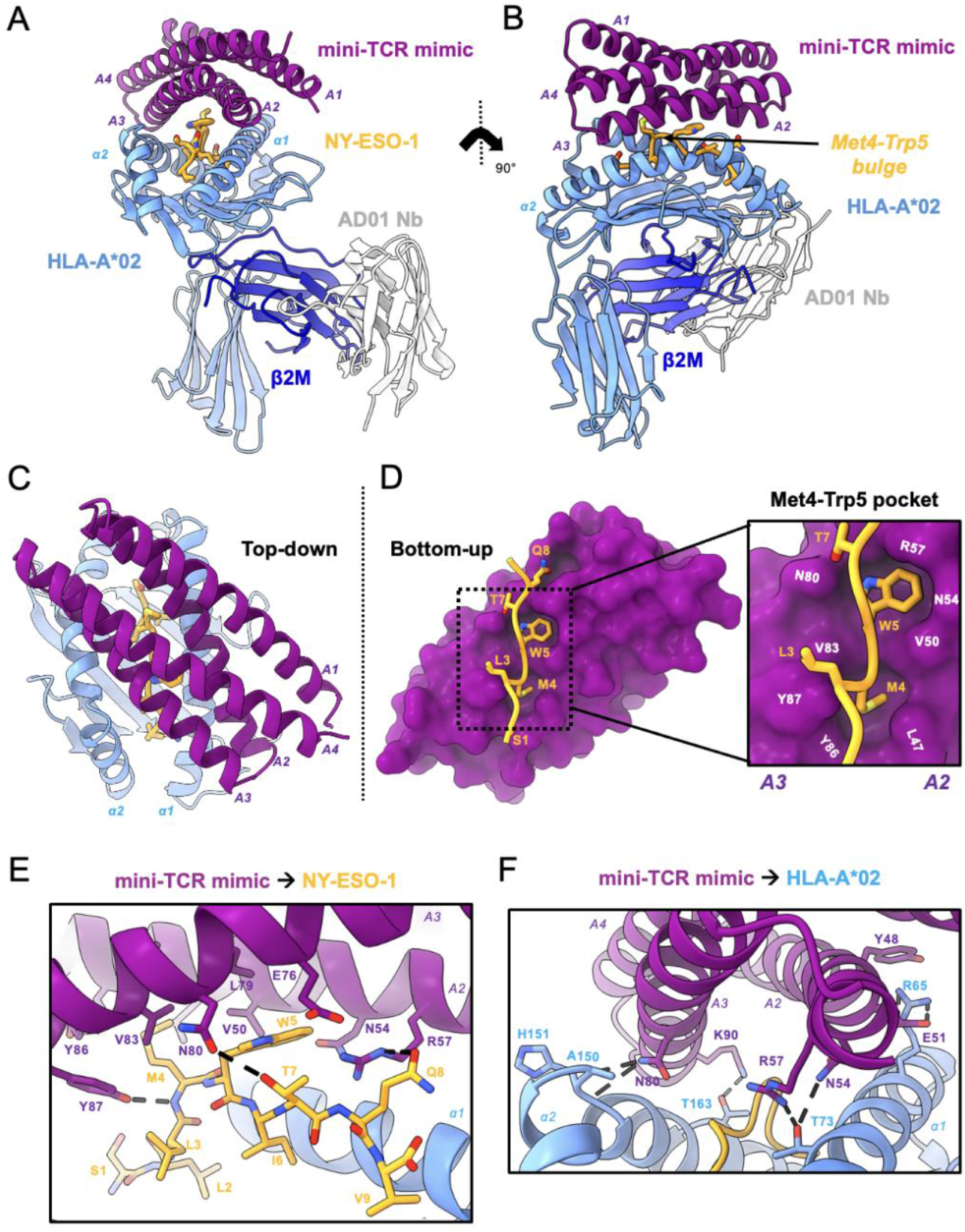
High-resolution crystal structure reveals peptide-specific interactions. **(A)** Front view of mini-TCRm/pMHC complex (PDB ID 9MIN). A1-A4 alpha-helices of mini-TCRm (purple); α1 and α2 helices of HLA-A*02 and full chain (light blue); NY-ESO-1 peptide (gold); β2M (dark blue); AD01 nanobody (white). **(B)** Side view of complex with Met4-Trp5 bulge on the NY-ESO-1 peptide. **(C)** Top-down view of complex. **(D)** Bottom-up view of peptide-specific interactions with surface of mini-TCRm (purple). Box zooming in on NY-ESO-1’s Met4-Trp5 motif (gold) fitting into mini-TCRm pockets formed between the A2 and A3 helices (key residue labels in white). **(E)** Hydrogen bonds (dashed lines) and key peptide-centric residues between mini-TCRm and NY-ESO-1 peptide. **(F)** Hydrogen bonds and salt bridges (dashed lines) between mini-TCRm and HLA-A*02.

Comparison with the AlphaFold prediction of the mini-TCRm/pMHC complex reveals that the computational model accurately predicts the diagonal docking orientation of the binder, including the hydrophobic pockets that accommodate the Met4-Trp5 bulge (R.M.S.D 0.410 Å for 128 Cα atoms) (fig. S1A). However, three subtle differences are observed between the prediction and the crystal structure, which affects peptide-specific contacts. First, AlphaFold predicts a hydrogen bond between Glu76 of the binder and Gln8 of NY-ESO-1, which is not present in the crystal structure (fig. S1B). Second, AlphaFold fails to predict an interaction between Asn80 and Thr7, yet a hydrogen bond is observed in the crystal structure (fig. S1C). Third, Arg57 makes a hydrogen bond with Gln8 in our structure, but AlphaFold predicts it to pair with the backbone of Ile6 instead (fig. S1D). Taken together, these discrepancies highlight areas where structural validation and structure-guided optimization add value to precise refinement of *de novo* TCR mimic specificity.

### Comparison to existing antibody TCR mimic and natural TCR

When comparing the mini-TCR mimic to an existing NY-ESO-1-specific Fab (3M4E5 Fab – PDB: 3HAE) and a natural TCR (1G4 TCR – PDB: 2BNQ) (*26, 35*), its compact and rigid α-helical structure is the most striking difference (Fig. 3A). The designed binder is less than half the height of the Fab or TCR, suggesting that it could achieve closer proximity between a T cell and target cell in a bispecific T cell engager (TCE) format. Unlike the Fab and TCR, which extend flexible loops into the MHC groove to engage the peptide, our binder primarily interacts with the peptide’s bulging, upward-facing side chains through a groove between its helices (Fig. 3B). While the Fab and TCR use their CDR loops to create a large pocket simultaneously accommodating Met4 and Trp5, the designed binder uses two tight shape-complementary pockets to engage each residue independently (Fig. 3C). In terms of docking footprints, the designed binder interacts with 19 MHC residues, compared to 14 for the Fab and 13 for the TCR. Nine of these residues—Arg65, Lys66, Ala69, Gln72, Thr73, Val76 on the α1 helix, and Lys146, His151, Gln155 on the α2 helix—are shared across all three footprints and are classical MHC “anchor residues” used by the germline-encoded CDR loops of natural TCRs (*20*). Overall, our mini-TCR mimic exhibits the largest total buried surface area (1207.7 Å^2^ for the TCR and 1055 Å^2^ for the Fab), but it buries the smallest proportion of peptide surface (36.5% for the TCR and 38.5% for the Fab). This is largely explained by the extended α-helices of the mini-TCR mimic, which enables more extensive contact across the top of the MHC. Despite this, our mini-TCR mimic maintains peptide specificity with the highest binding affinity (Fig. 3C), suggesting that it forms more focused peptide contacts with dispersed MHC interactions to stabilize the binding. Furthermore, its rigid α-helical structure reduces the entropic penalty of binding by limiting conformational flexibility, in contrast to the Fab and TCR. The increased conformational freedom of the Fab and TCR’s CDR loops might contribute to their unpredictable cross-reactivity profiles.

**Fig. 3.**
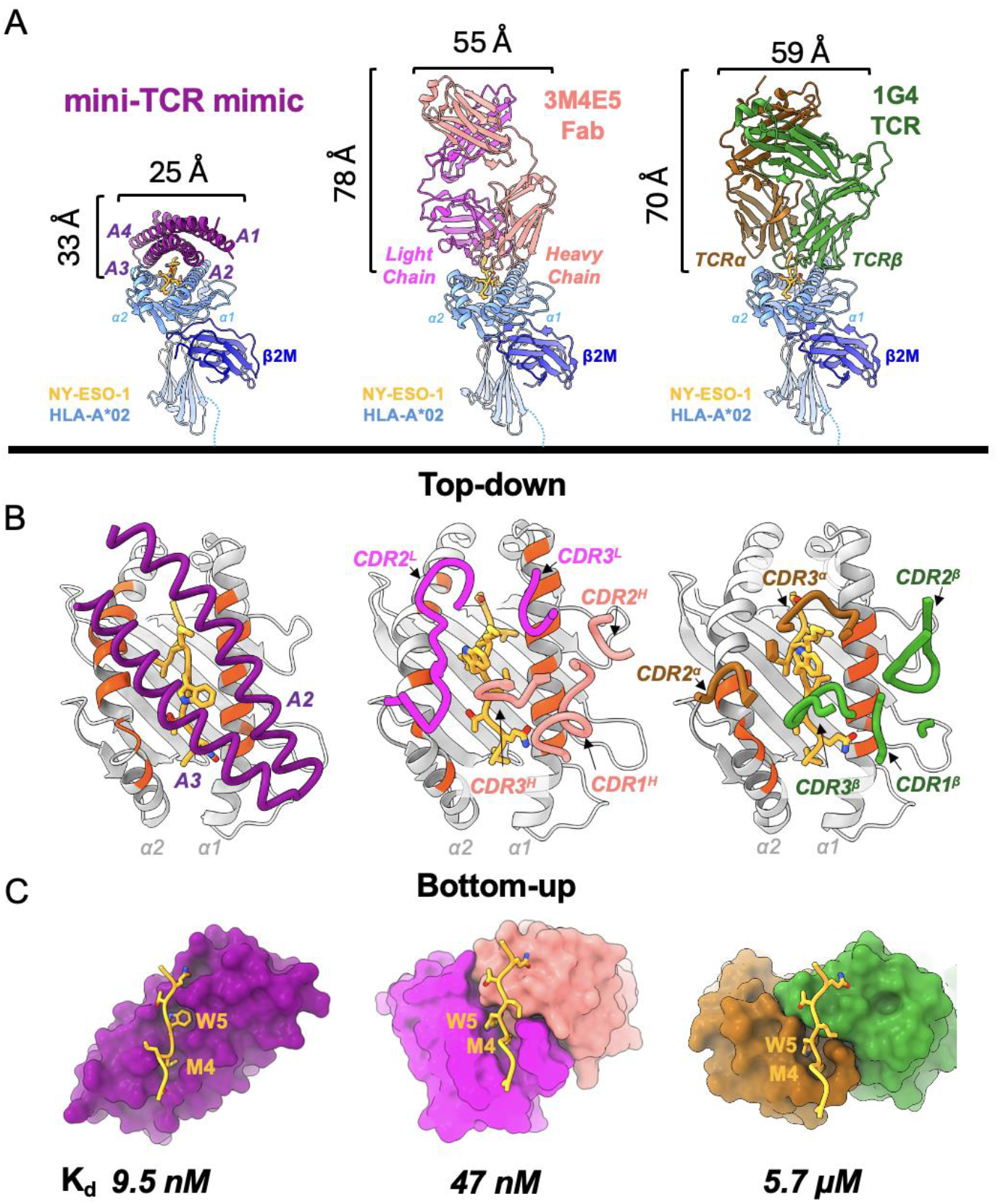
Comparison to existing antibody TCR mimic and natural TCR. **(A)** Front view comparison between mini-TCRm (purple), 3M4E5 Fab (magenta and salmon), and 1G4 TCR (brown and green) bound to NY-ESO-1 peptide (gold) and HLA-A*02 (light blue). Height and width of binders shown in Angstroms (Å). **(B)** Top-down comparison of docking footprints of each binder with MHC contact residues (red orange) and NY-ESO-1 peptide side chains (gold). **(C)** Bottom-up comparison of interactions between the NY-ESO-1 peptide (gold) and binding pockets, with the Met4-Trp5 motif labeled. Dissociation constants (K_d_) for each complex is shown at the bottom. nanomolar, nM; micromolar, μM.

### Structure-guided identification of off-target peptides from the human proteome

Using our crystal structure as a template, we sought to stress-test the out-of-the-box specificity of our TCR mimic. First, we needed to confirm the key peptide-specific hotspots observed in the structure, so we pulsed T2 cells with alanine-scanned variants of NY-ESO-1 and stained with our binder. Based on inspection of the crystal structure’s hydrophobic pockets, we hypothesized that chemically similar hydrophobic residues at positions 4 and 5 could also bind, so we included Leu4 and Phe5 mutants in the scan. We observed that mutation of either Met4 or Trp5 to alanine, completely abrogated binding; however, mutation to Leu4 or Phe5 rescues binding (Fig. 4A). This confirmed our hypothesis that the Met4-Trp5 bulge best stabilizes the complex, but also that chemically related residues can fit into the binder’s hydrophobic pockets. As expected, Thr7 and Gln8 mutations had minimal effect on binding when the Met4-Trp5 motif was present. Based on this information, we anticipated that off-target peptides with the highest chemical similarity to NY-ESO-1 at positions 1, 4, and 5 would be most likely to bind.

**Fig. 4.**
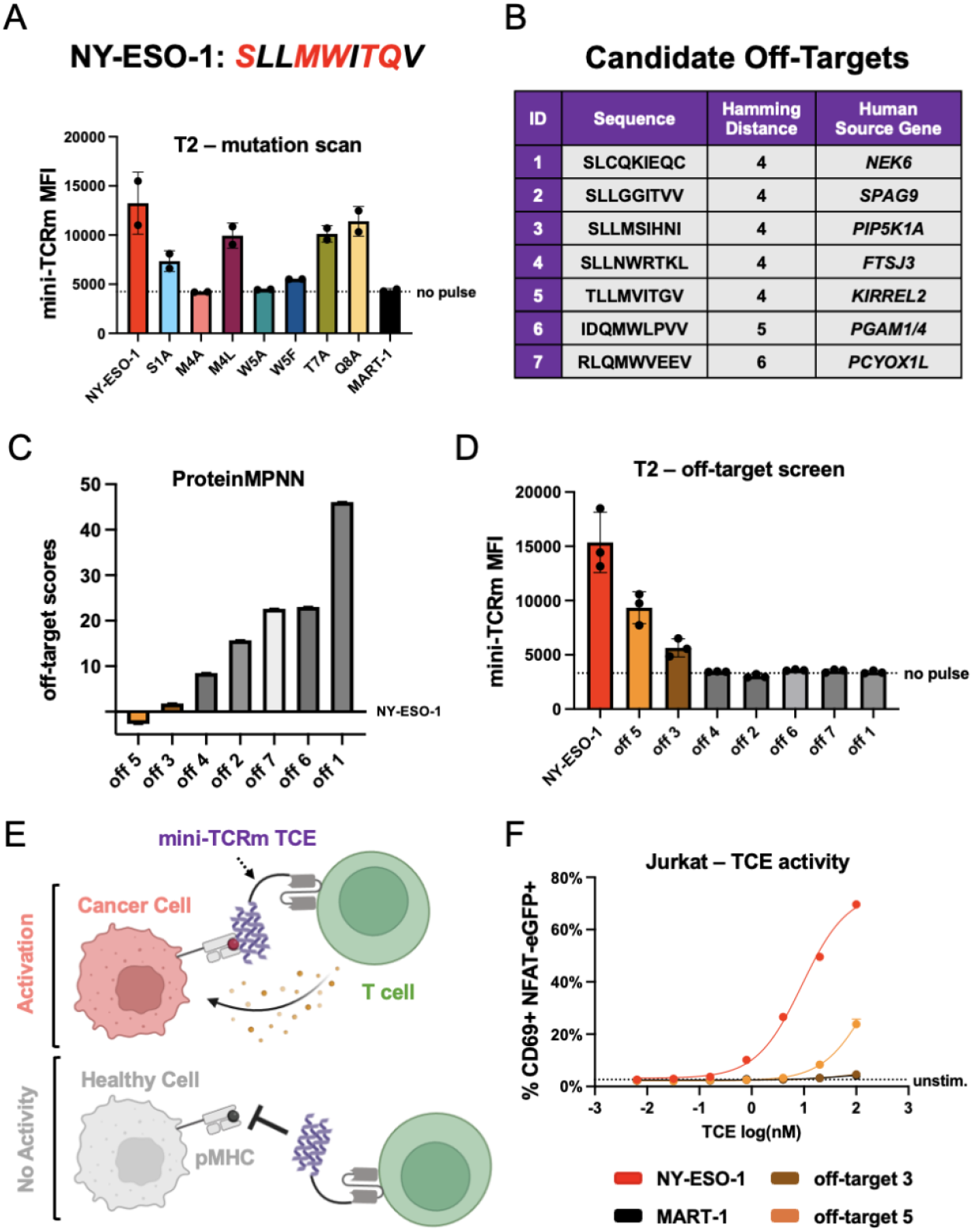
Structure-guided identification of off-target peptides reveals a robust therapeutic window. **(A)** Mutation scanning the NY-ESO-1 peptide (sequence shown above plot; mutated residues in red) and pulsing on T2 cells. Variants are named by original residue, position, followed by the mutation. Measured mini-TCRm MFI (Mean Fluorescence Intensity) by flow cytometry. **(B)** Table of candidate off-targets from the MHC Motif Atlas. **(C)** Ranking of candidate off-targets based on ProteinMPNN and the crystal structure. **(D)** Pulsing candidate off-target peptides on T2 cells and staining with mini-TCRm. **(E)** Schematic of a mini-TCRm T cell engager (TCE; purple) activating T cells against a cancer cell (salmon) with an on-target peptide (red) but not a healthy cell (gray) with an off-target peptide (black). **(F)** T2-Jurkat co-culture assay with pulsed peptides and mini-TCRm TCE. T cell activation shown as percentage of CD69+ NFAT-eGFP+ Jurkat cells versus no-pulse control from flow cytometry analysis. TCE dose shown in nanomolar on the log scale (log(nM)).

To test this hypothesis, we performed a simple Hamming distance search against the NY-ESO-1 peptide with 14,363 mass spectrometry (MS) detected 9-mer HLA-A*02 peptides from the MHC Motif Atlas (*39*). We restricted our search space to this dataset because not all plausible peptides from the human transcriptome are found at the cell surface due to differences in gene expression level, peptide competition, and processing. We reasoned that MS detection would provide a threshold for peptides found on the cell surface in sufficient quantity. After ranking the dataset by Hamming distance, we found only 5 out of 14,363 peptides had the lowest Hamming distance of 4 mutations away from NY-ESO-1 and derived from the human proteome (Fig. 4B). All other peptides were 5 or more mutations away from NY-ESO-1, which we believed would have a much lower likelihood of fitting our peptide’s target conformation and chemistry. To further sample these peptides, we performed another similarity search focused only on the Met4-Trp5 motif. This time, we found 2 out of 14,357 peptides with an exact match for the Met4-Trp5 motif, but overall Hamming distances of 5 and 6. These 7 peptides provided a simple and robust test set for off-target analysis. Next, we explored whether ProteinMPNN could rank these candidate peptides by their compatibility with the crystal structure. Providing the NY-ESO-1 peptide backbone as input and conditioning on the entire crystal structure, we used ProteinMPNN to assign an “off-target score” based on each peptide’s negative log-likelihood subtracted by the original NY-ESO-1 peptide’s negative log-likelihood. Intuitively, this approach predicts how well each off-target peptide fits the original peptide structure, conditioned on the presence of the mini-TCR mimic and MHC. We hypothesized that a peptide sequence with high likelihood to fit the structure would be a real off-target peptide. From this analysis, off-targets 3 and 5 exhibited the lowest off-target scores, closely aligning with NY-ESO-1, and therefore most highly predicted to bind (Fig. 4C). To experimentally confirm these predictions, we pulsed all peptides onto T2 cells and stained with our mini-TCR mimic. Consistent with the *in silico* analysis, off-targets 3 and 5 were true off-target peptides, though both stained more weakly than NY-ESO-1 (Fig. 4D). Off-target 3 originates from PIP5K1A, a ubiquitously expressed kinase, while off-target 5 derives from KIRREL2, a transmembrane cell adhesion molecule predominantly found in pancreatic beta cells. The AlphaFold-predicted structures of these off-targets revealed the retention of the Met4 residue, a hydrogen bond between Tyr87 and Met4’s backbone, and a small polar residue at position 1 to be the key binding features, with accommodation of peptide residues 5-8 subtly shifting the amount of buried surface area (fig. S2). Off-target 5 had the largest predicted BSA (1213.6 Å^2^) compared to off-target 3 (940.2 Å^2^), in line with our staining results. While these findings provide a compelling proof-of-concept for structure-guided off-target predictions, further large-scale validation is required to establish the accuracy and limitations of this approach.

### Peptide-specific T cell engagers provide a therapeutic window around off-targets

We then sought to determine whether our binder could function as a peptide-specific T cell engager (TCE) (Fig. 4E). We designed our TCE as a fusion between the anti-CD3ε scFv from blinatumomab (*40*) and our binder, connected by a short 5 amino acid Gly-Ser linker. Then, we pulsed T2 cells with NY-ESO-1, MART-1, off-target 3, and off-target 5 and co-cultured in a 1:1 ratio with a Jurkat reporter cell line that expresses eGFP after TCR signaling. A dilution series of our TCE was added and the proportion of activated CD69+ eGFP+ Jurkats was measured by flow cytometry the next day. As expected, in the presence of NY-ESO-1 but not MART-1 peptide, we measured potent T cell activation with an EC_50_ of 9.1 nM and nearly 70% of Jurkats activated with 100 nM engager (Fig. 4F). Surprisingly, no T cell activation was observed against off-target 3, despite our staining data. For off-target 5, we see ∼25% Jurkat activation with 100 nM engager but a log-shifted EC_50_ of 174 nM. However, because of the engager’s therapeutic window, as we drop the dose to 10 nM we retain potent NY-ESO-1 activity but zero off-target activity. Therefore, while we acknowledge that further optimization campaigns might reduce off-target specificity in terms of binding, our mini-TCR mimic might provide a starting point closer to an acceptable therapeutic window that will greatly reduce experimentation for human therapeutics.

## Discussion

Here, we have shown that an alternative structural scaffold to antibodies, with ideal protein engineering properties, offers a powerful platform for designing peptide-specific TCR mimic immunotherapies. By leveraging deep learning tools (*33, 34*), we have demonstrated that the time horizon to lead discovery of TCR mimics with robust peptide specificity and therapeutic window can be considerably shortened. While our current molecule might require further optimization and off-target screening to be deemed a clinical candidate, we believe it provides an initial starting point that could be closer to development than traditional antibody screening methods. Our approach could be particularly significant for personalized cancer therapies based on tumor antigens from exome sequencing (*1,2*), enabling real-time engineering of therapeutics in a clinically meaningful timeline. The α-helical platform also offers the advantage of modularity over traditional antibodies. One could envision these mini-TCR mimics engineered into multi-specific or multi-valent biologics with greater flexibility and complexity than currently feasible with the VH/VL heterodimer of antibodies, and potentially unlock new targeting strategies. Additionally, the relative rigidity of our α-helical scaffold provides significant computational advantages over traditional antibody approaches, as the free energy consequences of mutation are more accurately predicted compared to the flexible CDR loops of antibodies (*29-32*). With this restricted chemical and conformational search space, scaling up our *in silico* off-target screening approach could rapidly identify dangerous off-targets and if necessary, inform a subsequent round of structure-guided optimization. While the crystal structure was crucial for this work in validating and noting discrepancies in the design process, we anticipate that continued advancements in computational methods will progressively reduce the need for structural validation, such that it will not be necessary in future discovery efforts.

## Acknowledgments

We thank all members of the Garcia Lab for helpful discussions. We thank Jay Nix at the ALS for assistance with X-ray data collection. The Berkeley Center for Structural Biology is supported in part by the Howard Hughes Medical Institute. The ALS is a Department of Energy Office of Science User Facility under Contract No. DE-AC02-05CH11231. Figure 1A-C and Figure 4E were made with BioRender. We also thank V. Mallajosyula for valuable support with protein biochemistry.

## Funding

National Institute of Health grant 5R01AI103867-08 (K.C.G.).

National Cancer Institute grant OT2CA297242 (K.C.G.).

This work was delivered as part of the MATCHMAKERS team, supported by the Cancer Grand Challenges partnership financed by CRUK (CGCATF-2023/100006), the National Cancer Institute (OT2CA297242), and the Mark Foundation for Cancer Research (K.D.H., K.M.J., K.C.G.).

Yosemite Fund (K.C.G.).

K.C.G. is an investigator of the Howard Hughes Medical Institute (HHMI).

## Author contributions

K.D.H. and K.C.G. conceived the project and wrote the manuscript; K.D.H., X.X., K.M.J., A.D., M.O., and S.C.W. designed and performed experiments; K.D.H., X.X., K.M.J., S.C.W. determined crystal structures; X.C. and N.W. provided experimental reagents and protocols. All authors edited and approved the manuscript.

## Data and materials availability

The crystallographic model and integrated intensities have been deposited in the RCSB PDB with accession code 9MIN. Raw diffraction images have been deposited in the SBGrid databank. All other data are available in the manuscript or the supplementary materials.

## Supplementary Materials

Materials and Methods

Figs. S1 to S3

Table S1

References (*41*–*52*)

