## supplementary_materials for "*De novo* design and structure of a peptide-centric TCR mimic binding module"

### Materials and Methods

Cell culture

T2 (ATCC CRL-1992) and Jurkat NFAT-eGFP reporter cells were grown in RPMI 1640 Medium with 1x GlutaMax (Gibco) supplemented with 10% v/v fetal bovine serum (FBS), penicillin-streptomycin (Gibco) , MEM-NEAA (Gibco), HEPES (Gibco), Na Pyruvate (Gibco) and maintained at 37°C with 5% $CO_2$. Expi293F cells were grown in serum-free Expi293 expression media (Thermo) and maintained at 37°C with 5% $CO_2$. BL21 (DE3) *E. coli* (Novagen) were grown in LB media with carbenicillin or kanamycin at 37°C. EBY100 *S. cerevisiae* (ATCC MYA4941) were grown in SDCAA media pH 4.5 at 30°C.

Mini-TCR mimic design

RFdiffusion model weights and code were downloaded from the RFdiffusion GitHub repository (https://github.com/RosettaCommons/RFdiffusion). Backbone structures of four-helix α-helical bundles were sampled between 80-120 residues and a single optimal scaffold was selected by visual inspection for RFdiffusion fold conditioning. The target peptide-MHC structure for NY-ESO-1 HLA-A*02 was created with AlphaFold2. Hotspot residues on the peptide were selected to be Met4, Trp5, Thr7, and Gln8. ProteinMPNN and AlphaFold2 model weights and code were downloaded according to the dl_binder_design Github repository (https://github.com/nrbennet/dl_binder_design). Scaffolds generated by RFdiffusion were then fed into the dl_binder_design pipeline for sequence design and scoring by ProteinMPNN and AlphaFold2. All output designs were ranked from lowest to highest iPAE. Designs were considered hits if their iPAE was less than 10.0.

Bispecific T cell engager design

T cell engagers were designed as C-terminal fusions with mouse serum albumin (MSA) in the pD649 mammalian expression vector, containing an N-terminal hemagglutinin (HA) signal peptide and C-terminal 8x-His tag. The anti-human CD3ε scFv L2K-07 from blinatumomab (*40*) was fused C-terminally to MSA with a seven amino acid Gly-Ser linker, followed by a five amino acid Gly-Ser linker and the mini-TCR mimic.

Yeast display screening by flow cytometry

Gene blocks for the top mini-TCR mimic designs were ordered with overhangs for C-terminal display on digested pCT3CBN vector. Individual gene blocks and vector were mixed and electroporated as single clones into electrocompetent EBY100 yeast. Electroporation, rescue, expansion, and induction were performed as previously described (*41*). Data was collected on an Accuri C6 flow cytometer (Beckman Coulter). Mini-TCR mimics were stained with AlexaFluor-647-conjugated streptavidin (SA) control, 100 nM on-target tetramer (NY-ESO-1 HLA-A*02; Acro Biosystems), and 100 nM off-target tetramer (MART-1 HLA-A*02; Acro Biosystems). Designs that stained for NY-ESO-1 but not MART-1 were considered hits and advanced for further characterization.

Surface plasmon resonance

A BIAcore Control T100 (GE Healthcare) was used to measure $K_d$ by the multi-cycle kinetics method. Biotinylated pMHC (Acro Biosystems) was immobilized on a SA sensor chip (Cytiva) at 100-140 response units. Purified mini-TCR mimic was injected at varying concentrations (0 -156 nM) with 10 mM HEPES, 150 mM NaCl, 0.05% v/v Surfactant P20, pH 7.4 (HBS-P+) (Cytiva) for 60 s at a flow rate of 30 μL $min^{-1}$, then dissociation was measured for 60 s with buffer flow. The signals of reference cells were subtracted from measurements. Data analysis was performed with BIAcore T100 evaluation software.

Protein production and purification

For mini-TCR mimic production, gene blocks were cloned into pETDuet1 vectors with a C-terminal 6x-His tag. Plasmids were then transformed into competent BL21(DE3) *E. coli* and allowed to grow overnight shaking at 37°C in LB starter cultures. The next day, starter culture was added to 1 L of LB media and continued to incubate until OD = 0.6-0.8. Culture was then induced with 0.2 mM isopropyl β-D-1-thiogalactopyranoside (IPTG) (Sigma Aldrich) overnight shaking at 18°C. Cells were then pelleted and lysed with B-PER Bacterial Protein Extraction Reagent lysis buffer (Thermo) according to the manufacturer's protocol and protein was purified with Pierce Nickel-NTA resin (Thermo) followed by SEC on a Superdex 200 column (Cytiva) in 20 mM HEPES pH 7.4, 150 mM NaCl (HBS).

For T cell engager production, designed plasmids were transiently transfected into the Expi293F mammalian cell line using Expifectamine transfection reagent. Cell supernatant was harvested and purified 96 to 120 hours later with Nickel-NTA resin followed by SEC on a Superdex 200 column in HBS.

For anti-β2M nanobody production, a gene block of nanobody AD01 (*41*) was cloned into pD649 with an N-terminal HA signal peptide and C-terminal 8x-His tag. The plasmid was transiently transfected into Expi293F cells and after five days, the supernatant was collected and the protein was batch purified on Ni-NTA resin, followed by SEC on a Superdex 75 column (Cytiva) in HBS. AD01 was stored in flash-frozen aliquots before use.

For production of HLA-A*02 and β2M inclusion bodies, gene blocks were cloned into pETDuet1 vectors with no tags. Plasmids were transformed into competent BL21 BL21(DE3) *E. coli* and allowed to grow overnight shaking at 37°C in LB starter cultures. The next day, starter culture was added to 1 L of LB media and continued to incubate until OD = 0.5-0.7. Culture was then induced with 1 mM IPTG shaking at 30°C for 3 hours. Cells were pelleted and resuspended in resuspension buffer (50 mM Tris-HCl pH 8.0, 1 mM EDTA, 10 mM DTT). Cell pellets were then lysed with lysis buffer (50 mM Tris-HCl pH 8.0, 1% Triton X-100, 100 mM NaCl, 10 mM DTT) rotating for 20 minutes at room temperature. Lysate was sonicated and pelleted. Inclusion body pellets were then washed and pelleted three times with detergent-based wash buffer (50 mM Tris-HCl pH 8.0, 0.5% Triton X-100, 1 mM DTT, 100 mM NaCl, 1 mM EDTA). Finally, the preparation was washed with detergent-free wash buffer (50 mM Tris-HCl 8.0, 1 mM EDTA, 1 mM DTT, 0.2 mM PMSF) and pelleted. Inclusion bodies were solubilized with urea buffer (8M urea, 20 mM Tris-HCl pH 8.0, 0.5 mM EDTA, 1 mM DTT) and frozen at -80°C until use.

Refolding and purifying peptide-MHC

Peptides for refolding were synthesized by Elim Biopharm. Refolding buffer was prepared stirring at 4°C (100 mM Tris-HCl pH 8.0, 2 mM Na EDTA, 400 mM L-Arginine-HCl, 0.5 mM oxidized glutathione, 5 mM reduced glutathione) with 10 mg of NY-ESO-$1_{157-165\ (C9V)}$ peptide. 10 mg of each solubilized inclusion body was combined and slowly added to refolding buffer (*43*). The refolding mixture was then dialyzed four times against 10 mM Tris-HCl buffer. Refolded pMHC was purified by SEC Superdex 200, followed by MonoQ (GE Healthcare) columns.

<u>X-ray crystallography of complex</u>

Mini-TCR mimic, refolded NY-ESO-1 pMHC, and AD01 nanobody were complexed in a 3:1:1 molar ratio and incubated at 4°C overnight with 1:1000 (w/w) carboxypeptidases A and B. The complex was purified by a SEC Superdex 200 column and the co-eluting fractions were confirmed by SDS-Page gel. Purified complex was concentrated to 10.8 mg/mL and crystallized using the Index screen (Hampton Research) in 0.2 M ammonium acetate, 100 mM bis-tris pH 5.5, and 25% PEG 3350. Crystals were cryoprotected by addition of 30% glycerol and flash cooled in liquid nitrogen. Diffraction data were collected at Advanced Light Source (ALS) beamline 8.2.1. Initial data processing with XDS and pointless (*44*, *45*) suggested the space group $P2_12_12$ with cell dimensions a = 95.4 Å, b = 98.5 Å, c = 77 Å. Structure solution by molecular replacement using Phaser (*46*) with AlphaFold2 models (*37*) of the mini-TCR mimic and the extracted crystal structure of NY-ESO-1 pMHC (PDB: 3HAE). One copy of the complex was found in the asymmetric unit, but the electron density maps were barely interpretable. Reinspection of the diffraction data revealed a true space group of $P2_12_12_1$ with a doubled c axis of 155.7 Å. A translational noncrystallographic symmetry vector parallel to the c-axis was found to have caused in very weak reflections for l = 2n. Repeating the molecular replacement procedure identified two complexes in the asymmetric unit and produced interpretable electron density maps (fig. S3). The model was built and refined in iterative cycles of interactive and automated refinement using Coot (*47*) and Phenix (*48*) using torsional NCS restraints (*49*). TLS parameters were assigned using TLSmd (*50*). The final structure had 98.71% of residues in the favored region of the Ramachandran plot, with 0 outliers. Crystallographic data and refinement statistics are reported in Table S1. Crystallographic software for this project was installed and configured using SBGrid (*51*). PDBePISA software was used to calculate buried surface area for the mini-TCR mimic, 3M4E5 Fab, and 1G4 TCR and to confirm, count, and compare key interactions observed in the crystal structures (*52*).

<u>T2 peptide pulsing assays</u>

Peptides for pulsing assays were ordered from Elim Biopharm. 0.5E6 T2 cells per well were plated into a 96-well plate and 100 μM peptide was prepared in serum-free RPMI 1640 with 1x Glutamax (Gibco). Peptides were added to T2 and allowed to incubate at 37°C with 5% $CO_2$ for 2 hours. Excess peptide was washed two times with serum-free media. Pulsed cells were then stained with 1 μM mini-TCR mimic in PBS for 20 minutes on ice, followed by washing and staining with FITC-conjugated anti-His tag antibody (50:1, Biolegend) for 20 minutes on ice. Plate was washed and the mean fluorescence intensity (MFI) was acquired using a CytoFlex flow cytometer (Beckman Coulter).

<u>T cell engager signaling assays</u>

T2 cells were pulsed with 10 μM peptide for 2 hours as previously described. 0.04E6 T2 and 0.04E6 Jurkat NFAT-eGFP per well were added to a 96-well plate. Starting from 100 nM mini-TCRm engager, seven 5:1 serial dilutions were added to each well, including controls with no engager. Co-cultures were incubated for 16-18 hours at 37°C with 5% $CO_2$. The next day, cells were washed and stained for 20 minutes on ice with Zombie Violet Live/Dead (500:1,

Biolegend), PE-conjugated CD3 antibody (50:1, UCHT1 clone, Biolegend), and APC/Cy7-conjugated anti-CD69 antibody (50:1, Biolegend). Data were collected on a CytoFlex flow cytometer with compensation applied. Jurkat NFAT-eGFP cells were identified as the $SSC_{lo}$ Live+ CD3+ population. Compared to controls with either no peptide pulsing or no engager, the percentage of these cells that were CD69+ NFAT-eGFP+ were considered activated Jurkat cells.

Selecting off-target peptides by Hamming distance

We downloaded all 9mer HLA-A*02 peptides from the MHC Motif Atlas and used a short Python script that calculates the Hamming distance for each peptide (compared to the natural NY-ESO-1 peptide – SLLMWITQC) to output a CSV file ranking this dataset by lowest to highest Hamming distance. This yielded six peptides with the lowest Hamming distance of four. Five of the six peptides were found in the human proteome and therefore included in the dataset. Next, we calculated the Hamming distance for peptides with the Met-Trp motif at positions 4 and 5 and sorted the dataset. This returned only two peptides with a Hamming distance of 0. Both were found in the human proteome and were therefore added to our final dataset.

Scoring off-target peptides with ProteinMPNN

Using our crystal structure, we generated a positional matrix of ProteinMPNN negative log-likelihood values for all twenty amino acids at each position in the peptide backbone. For each off-target peptide's amino acid sequence, we calculated the sum of negative log-likelihood values at each residue position. We performed ten replicates for each off-target peptide and averaged these runs. Then, we repeated this procedure for the original NY-ESO-1 peptide sequence and subtracted this value from each off-target peptide's score. This final value is what we term the "off-target score". Off-target scores less than 0 indicate that the new peptide fits the crystal structure backbone better than NY-ESO-1, whereas a score greater than 0 indicates a worse fit. Peptides were ranked by lowest to highest off-target scores

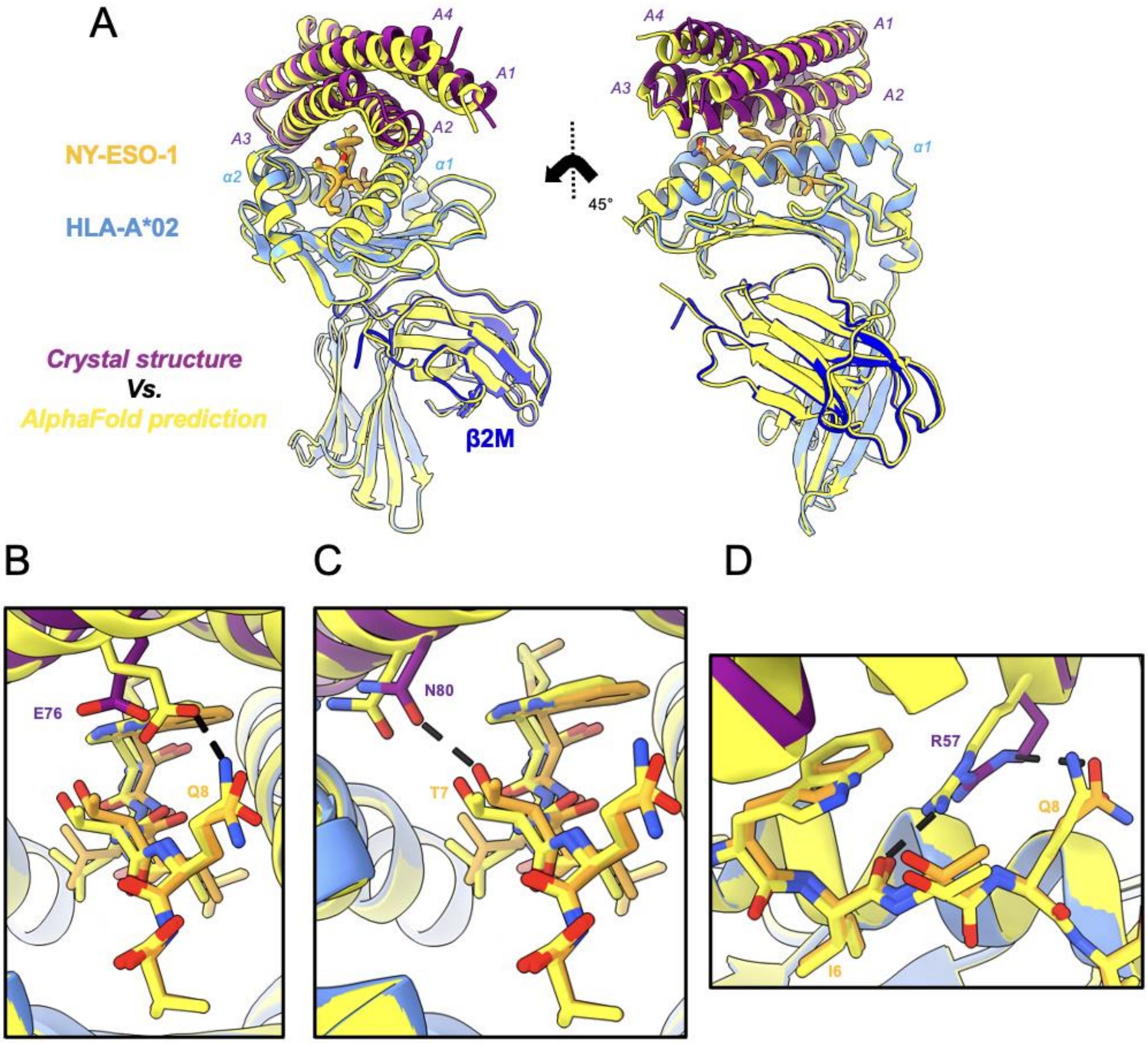


**Fig. S1. Comparison of crystallographic peptide contacts to the AlphaFold prediction.**

**(A)** Front and side view of the crystal structure aligned with the AlphaFold prediction (yellow). **(B)** Hydrogen bond in AlphaFold prediction between Glu76 of mini-TCRm and Gln8 of NY-ESO-1 (yellow). No interaction in crystal structure (purple and gold). **(C)** Hydrogen bond in crystal structure between Asn80 of mini-TCRm (purple) and Thr7 of NY-ESO-1 (gold). No interaction in AlphaFold prediction (yellow). **(D)** Hydrogen bond in crystal structure between Arg57 on mini-TCRm (purple) and Gln8 on NY-ESO-1 (gold). AlphaFold prediction makes a hydrogen bond between Arg57 and Ile6's backbone (yellow).

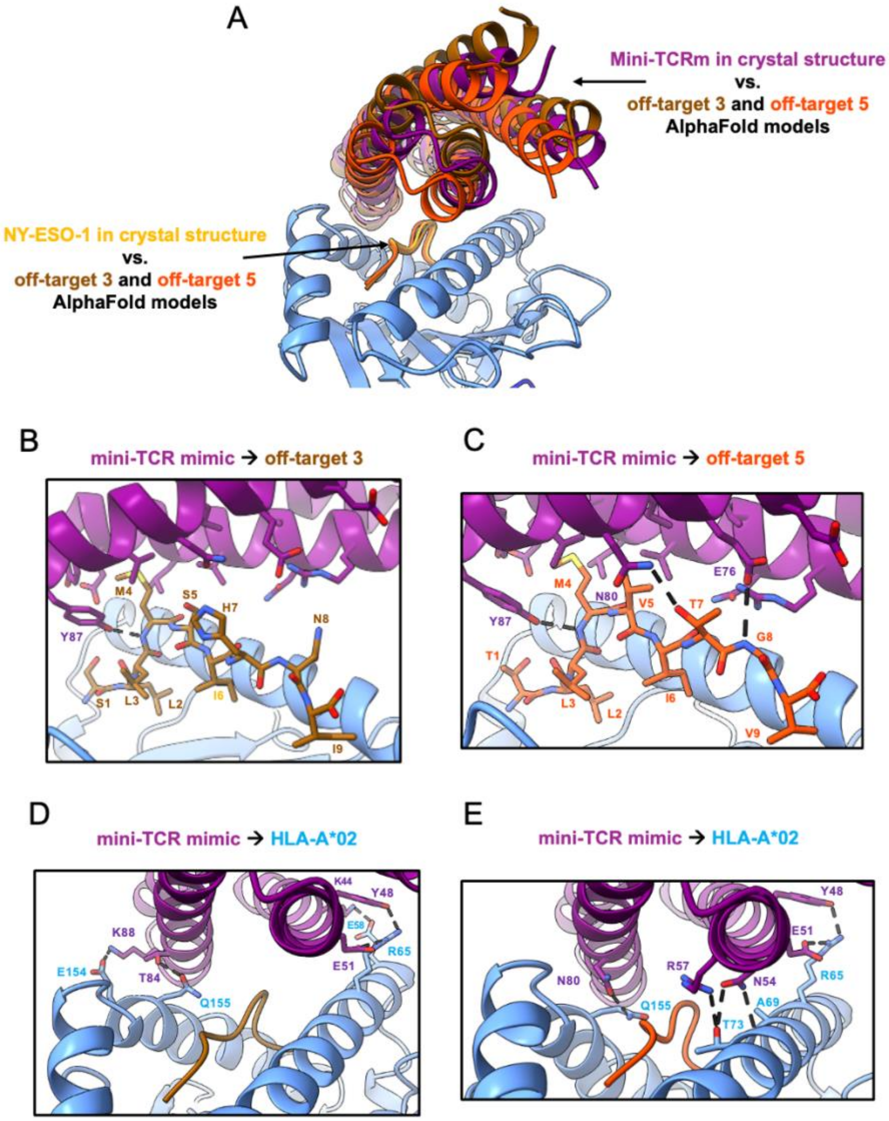


**Fig. S2. Comparison of the crystal structure to off-target AlphaFold predictions.**

**(A)** Alignment of mini-TCRm in the crystal structure (purple) docked over the NY-ESO-1 peptide (gold), versus the predicted docking geometry of the mini-TCRm over off-target 3 peptide (brown) and off-target 5 (orange). **(B)** Hydrogen bond in AlphaFold prediction between Tyr87 of mini-TCRm (purple) and the Met4 backbone of the off-target 3 peptide (brown). **(C)** Hydrogen bonds in AlphaFold prediction between Tyr87, Asn80, and Glu76 of mini-TCRm (purple) and Met4 backbone, Thr7, and Gly8 backbone of the off-target 5 peptide (orange). **(D)** Hydrogen bonds to HLA-A*02 in the AlphaFold prediction for off-target 3 (brown). **(E)** Hydrogen bonds to HLA-A*02 in the AlphaFold prediction for off-target 5 (orange).

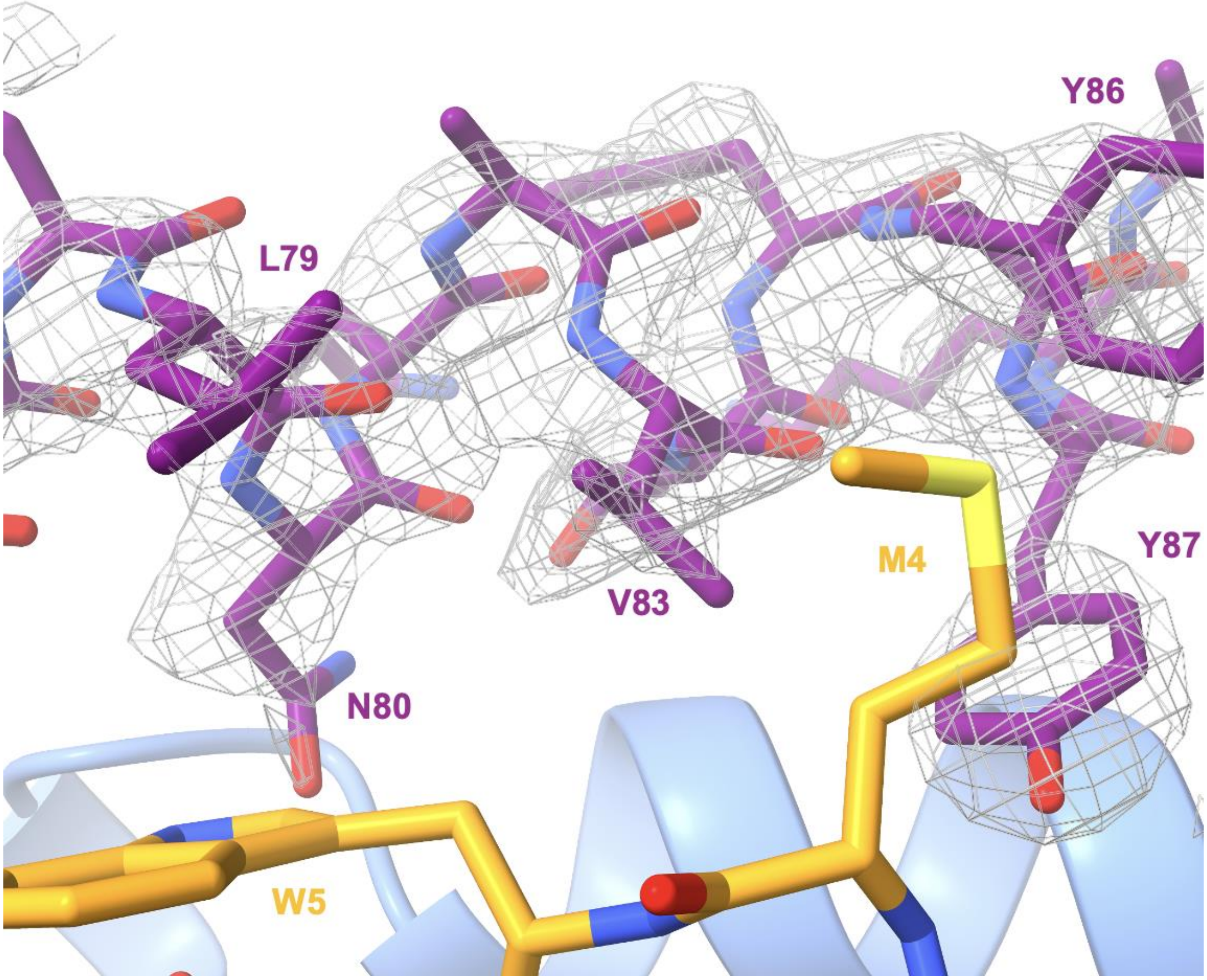


**Fig. S3. Fitting of the mini-TCRm/pMHC complex structure into the electron density map.** 2mFo-DFc electron density map contoured at 1σ (gray) around mini-TCRm A3 helix (purple). Key interacting mini-TCRm residues are labeled. NY-ESO-1 positions Met4 and Trp5 (gold) and the α2 helix of HLA-A*02:01 (light blue) are shown as sticks and cartoon, respectively.

**Table S1. Crystallographic Data and Refinement Statistics.**

| | Mini-TCR mimic + NY-ESO-1 + A*02:01 + AD-01 Nb |
|---|---|
| Wavelength (Å) | 1.00003 |
| Resolution range (Å) | 49.27 - 2.05 (2.12 - 2.05) |
| Space group | P 21 21 21 |
| Unit cell (Å, °) | 95.452 98.536 155.743 90 90 90 |
| Total reflections | 2159405 (218759) |
| Unique reflections | 92563 (9127) |
| Multiplicity | 23.3 (24.0) |
| Completeness (%) | 99.17 (100.00) |
| Mean I/sigma(I) | 8.49 (1.45) |
| Wilson B-factor ($Å^2$) | 33.17 |
| R-merge | 0.3252 (3.471) |
| R-meas | 0.3326 (3.545) |
| R-pim | 0.0686 (0.7168) |
| CC1/2 | 0.999 (0.349) |
| Reflections used in refinement | 91895 (9127) |
| Reflections used for R-free | 1384 (133) |
| R-work | 0.2549 (0.3981) |
| R-free | 0.2672 (0.4134) |
| Number of non-hydrogen atoms | 10482 |
| macromolecules | 10116 |
| ligands | 96 |
| solvent | 270 |
| Protein residues | 1258 |
| RMS(bonds) (Å) | 0.003 |
| RMS(angles) (°) | 0.53 |
| Ramachandran favored (%) | 98.55 |
| Ramachandran outliers (%) | 0.00 |
| Rotamer outliers (%) | 2.43 |
| Clashscore | 4.95 |

| | |
|---|---|
| **Average B-factor (Å$^2$)** | 60.86 |
| **macromolecules** | 61.41 |
| **ligands** | 54.57 |
| **solvent** | 42.61 |

Statistics for the highest-resolution shell are shown in parentheses.
